# Temporal dynamics of inflammatory and transcriptome changes in the myocardium in a murine model of cardiac arrest

**DOI:** 10.1101/2024.10.25.620238

**Authors:** Soumya Panigrahi, Alice Jiang, Angela Enriquez, Sanjana Tummala, Donald Rempinski, Kenneth E. Remy, Cody A. Rutledge

## Abstract

**Background:** It is increasingly recognized that inflammatory changes play a significant role in myocardial injury following cardiac arrest. However, the pathophysiological mechanisms responsible for functional changes in the myocardium following the event remain incompletely understood. Here, we characterize the transcriptomic and immunological changes in a mouse model of cardiac arrest to explore the roles of epinephrine and ischemia/reperfusion on myocardial inflammation.

**Methods and Results:** Male and female mice (C57BL/6J) were divided into three groups: (1) Naïve (anesthesia only), (2) Sham (anesthesia + saline injection + epinephrine), and (3) Arrest (8-minute cardiac arrest followed by resuscitation (anesthesia + potassium chloride + epinephrine). We monitored the animals over 0.5, 1, 3, and 7 days with serial echocardiography. Hearts were harvested for transcriptomic analysis and inflammatory changes as assessed by flow cytometry and immunohistochemistry. Plasma samples were collected for cytokine signaling analysis by multiplex ELISA. Our data indicate that the cardiac arrest group have increased infiltrating neutrophils in the myocardium and elevated plasma levels of pro-inflammatory cytokines, peaking at 0.5 days after the initial insult and subsequently resolving. In contrast, Sham mice displayed less pronounced inflammatory changes, which peaked at 1 day after the procedure. Notably, these inflammatory alterations coincided with cardiac stunning, as demonstrated by echocardiography.

**Conclusions:** In our murine model of cardiac arrest, epinephrine exposure and ischemia/reperfusion induce significant inflammatory changes in the myocardium with increased infiltrating neutrophils and elevated circulating cytokines which subsequently resolve in a time-dependent manner. These changes correlate with improving cardiac function by echocardiography.

## Introduction

Each year, over 356,000 people in the United States are treated for out-of-hospital cardiac arrests (CA)^1^. Unfortunately, the survival rate for these incidents remains low, with less than 10% of the patients surviving long-term^2^. Despite progress made in developing advanced emergency responses and the availability of prompt cardiopulmonary resuscitation (CPR), the challenges of managing CA in healthcare systems remain high^1,3,4^. While rapid resuscitation from CA improves outcomes, all survivors of CA undergo a milieu of molecular and metabolic deregulations systemically that cause pro-inflammatory immunological injury^5–7^. There currently exist no widely adopted therapeutics that significantly alter the poor outcomes of CA.

CA triggers a wide cascade of events that impact a number of tissues in the body, including the myocardium^8^. During CA, tissues suffer from severe oxygen and nutrient deprivation, leading to ischemia, hypoxia, ATP depletion, and accumulation of metabolic waste products, causing cellular acidosis^6^. Upon resuscitation, the sudden influx of oxygen generates reactive oxygen species (ROS), inducing oxidative stress and damaging cellular structures and exacerbating myocardial injury^9^. Disrupted calcium balance during ischemia leads to intracellular calcium overload upon reperfusion, impairing mitochondrial function, reducing ATP production, and eventually triggering cell death^10^. The ischemia-reperfusion process activates inflammation with neutrophils infiltrating the tissue and pro-inflammatory cytokines being released, further damaging the myocardium^11^. Collectively, oxidative stress, calcium overload, and inflammation lead to eventual cell death through apoptosis and necrosis, impairing the heart’s contractile function. Surviving cardiomyocytes may temporarily exhibit contractile dysfunction due to oxidative and metabolic changes, a condition known as myocardial stunning.

Myocardial stunning is a temporary state of mechanical cardiac dysfunction that occurs in heart tissue without necrosis after a brief interruption in blood flow, even when normal coronary blood flow is promptly restored^12,13^. A wide number of physiologic changes have been implicated in the pathogenesis of myocardial stunning including altered regional blood flow^14^, increased reactive oxygen species^9^, altered calcium handling and sensitivity^10^, and more recently, immune activation^15^. Systemic inflammation is also a contributor to post-cardiac arrest syndrome (PCAS), where increased circulating inflammatory markers are associated with worse functional outcomes^11,16^. Infiltrating immune cells are major contributors to cardiac response to ischemic injury and have been well characterized in myocardial infarction^17^.

While extensive studies have evaluated circulating markers of inflammation following cardiac arrest, little work has been done to evaluate transcriptomic changes of the myocardium following CA and to characterize the infiltrating immune cells into the cardiac tissues. In this study, we investigate the time-sensitive immune changes in the myocardium after CA using a murine model^18^. We hypothesize that the ischemia/reperfusion injury of CA leads to immune cell infiltration and cytokine dysregulation, leading to the development of a pro-inflammatory microenvironment in the affected myocardium. In this study, we focus on the time-dependent modulations of immune cells and the global transcriptome in cardiac tissue after CA. The intended outcome of this was to understand the temporal dynamics of immune cell activation in the affected myocardial tissues after global ischemia in the absence of coronary artery disease. Through this research, we aimed to gain insights into the inflammatory and immune responses triggered by cardiac arrest, which could inform potential therapeutic strategies and affect clinical outcomes.

## Methods

### Cardiac Arrest Model

Eight- to ten-week-old male and female C57BL/6J mice (Jackson Lab, Bar Harbor, ME, #000664) were randomly placed into either Naïve, Sham, or Arrest groups. Naïve mice were anesthetized using 5% isoflurane (Henry Schein, Melville, NY, #1182097) and euthanized prior to tissue collection. Sham and Arrest mice were treated with direct injection of saline or potassium chloride into the left ventricle (LV), by percutaneous, ultrasound-guided needle injection (Figure 1), as we have previously reported^18^. After anesthesia and intubation, 40 µL of saline or 0.5M KCl was injected. KCl-induced asystole was confirmed by ECG and doppler flow, lasting 8 minutes. At 8 minutes, 500 µL of epinephrine was injected, and chest compressions were initiated. Mice without return of spontaneous circulation (ROSC) within 3 minutes were euthanized. Arrest mice continued on ventilation until spontaneous breathing resumed and were then moved to recovery under a heat lamp for 2 hours.

**Figure 1:**
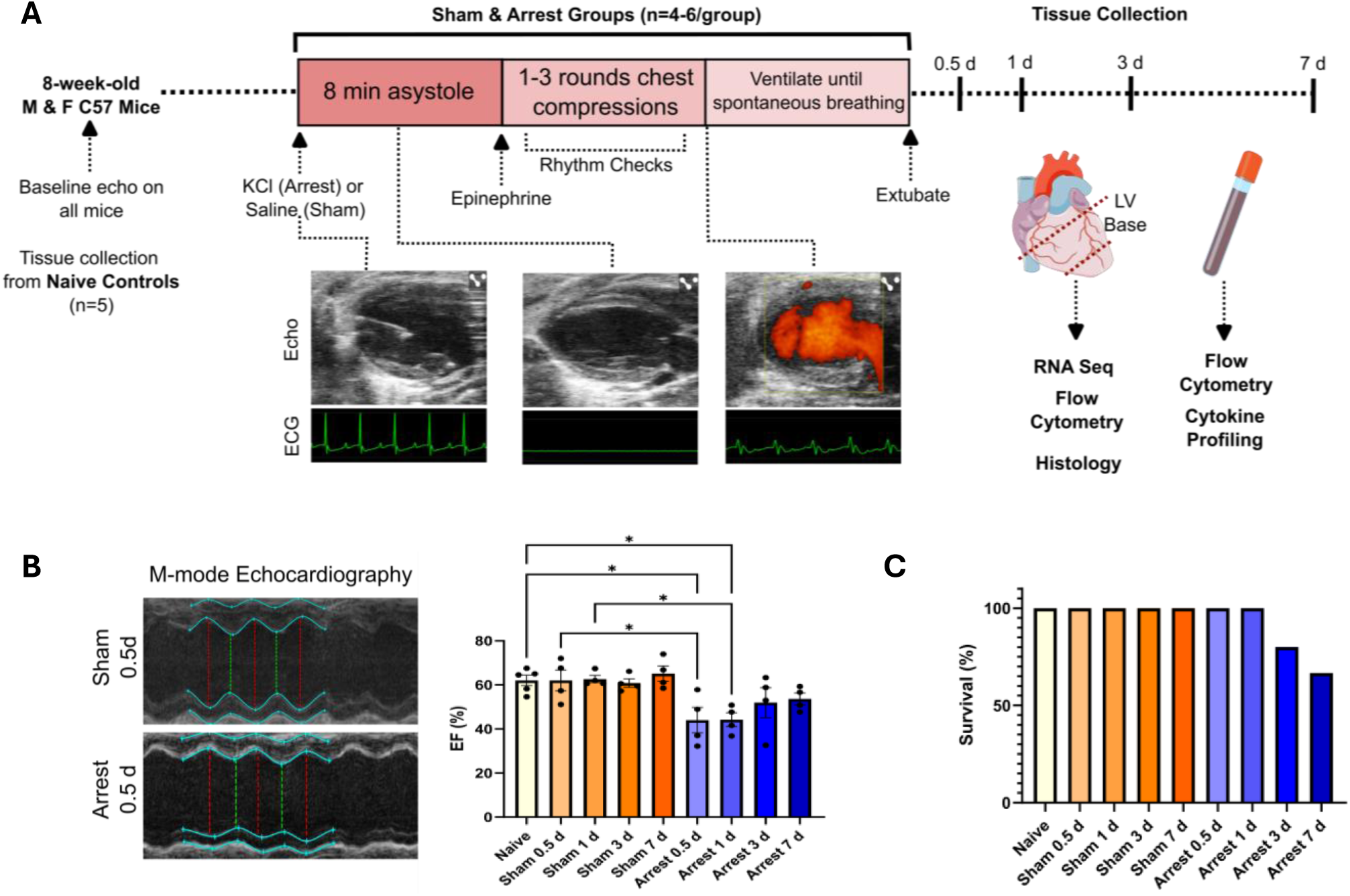
Mouse model of cardiac arrest. A. Study overview depicting potassium chloride injection to cause asystole with representative ECG data during normal sinus rhythm, asystole, and resuscitation, as well as description of end-points for tissue collection. B. Representative M- mode echocardiography (left panel) in Sham and Arrest (0.5d after procedure). Ejection Fraction (EF) in Naïve, Sham, and Arrest groups at study end points (right). C. Percent of all animals surviving to study endpoint. n=4-5/group. *=p <0.05 by ANOVA with Dunnett’s multiple comparison test.

### Treatment Duration, Echocardiography, and Tissue Collection

Baseline systolic function was assessed in Naïve, Sham, and Arrest mice using transthoracic echocardiography (Vevo 3100, Visual Sonics, Toronto, Canada), as previously described^19^. Heart rate was maintained at 400-500 bpm, and ejection fraction (EF) was calculated. Naïve mice had tissues collected immediately post-imaging, while Sham and Arrest groups underwent needle injection and recovered for 0.5, 1, 3, or 7 days. At least 4 mice per group were sampled at each time point, with additional mice included at 3 and 7 days to account for mortality. After recovery, mice were euthanized, and blood was collected via cardiac puncture. Blood was centrifuged, and serum was flash-frozen. Hearts were flushed with cold saline, and the apex was discarded to minimize local injury effects. The heart base was split; half was flash-frozen for analysis, and half was digested for flow cytometry. All mouse studies were performed at the Pittsburgh VA Medical Center (IAUCUC Protocol #1680736).

### Transcriptomics data and Immunoprofiling

RNA was extracted from 50 mg of frozen LV tissue using the RNeasy Mini Kit (Qiagen, Hilden, Germany). Concentration and integrity were measured with NanoDrop (Thermo Fisher, Waltham, MA) and TapeStation (Agilent Technologies, Santa Clara, CA), including samples with a 260/280 ratio ≥1.8 and RIN ≥6. Bulk RNA sequencing was performed by Novogene (Beijing, China) on an Illumina NovaSeq PE150 platform. Raw reads were quality-checked with FastQC and mapped to the reference mouse genome using HISAT2. One 3-day Sham sample was excluded due to poor quality. Transcript abundance was quantified with HISAT2 v2.0.5, and differentially expressed genes (adjusted p-value ≤ 0.05, log2(fold change) > 0) were identified with DESeq2. Reactome pathway analysis identified significant pathways (adjusted p-value < 0.05), and bulk RNA levels were cross-referenced with immune cell gene markers for immunoprofiling^20^ (Supplemental Table 2). Z-scoring was completed for each of the 16 immune cell populations and compared across all treatment groups as previously described^21^. Heat maps were generated using Heatmapper^22^ and Principal Component Analysis (PCA) plots were generated using ClutsVis^23^.

### Flow Cytometry

The heart base was minced in ice-cold phosphate-buffered saline, then incubated in 1 mL Dulbecco’s Modified Eagle Medium with 450 U/mL collagenase 1 (Sigma, #C0130), 60 U/mL hyaluronidase type 1-S (Sigma, H3506), and 60 U/mL of DNAse-1 (Sigma, #D4513) at 37°C for 1 hour. Tissue was homogenized, filtered through a 40 µm strainer into cold HBSS, and centrifuged at 400 g for 5 minutes. After ammonium-chloride potassium (ACK, Lonza, #10-546) lysis, cells were washed with HBSS, recentrifuged, and resuspended in FACS buffer. Samples were stained with antibodies (Supplemental Table 1) for 30 min at 4°C before flow cytometry. Spleens were similarly processed without enzymatic digestion. Whole blood underwent ACK lysis, washing, and antibody staining. Flow cytometry was performed on a Cytek Aurora and analyzed with FCS Express 7 (Denovo Software, Pasadena, CA; Supplemental Figures 3-5).

### Histology

Excised hearts were dissected by cutting through the ventricular tissue near the base of the heart. The cephalic edges of the ventricles were fixed in formalin and paraffin-embedded. 8 µM sections of LV were cut by microtome and mounted onto slides by the University of Pittsburgh Histology Core. Slides were incubated with goat anti-human/mouse myeloperoxidase (MPO; R&D Systems #AF3667, Minneapolis, MN) at 1 ug/ml for 1 hour at room temperature followed by incubation with anti-goat IgG HRP (R&D Systems #HAF109) and stained using DAB (brown) and counterstained with hematoxylin (blue). 40x images of the endocardium were obtained on an EVOS 5000 (ThermoFisher) and the number of neutrophils per high-powered field (HPF) was counted by a blinded observer using ImageJ (National Institute of Health).

### Cytokine analysis

Cytokine analysis was performed on serum samples using custom-made multi-plex cytokine plates (IFNg, IL-2, IL-6, TFN-α; IL-1β, IL17, GM-CSF, and CCL3) and analyzed on the ELLA microfluidic immunoassay system (ProteinSimple, San Jose, CA). 25 µL of plasma from each animal was diluted 1:1 with sample diluent and briefly vortexed. Each sample was loaded onto a sample inlet on an ELLA cartridge assay and wash buffer was loaded per the manufacturer’s recommendation. Sample results were using Simple Plex Runner v.3.7.2.0.

### Statistical Analysis

Data are presented as mean ± standard error. Significance was set at p ≤ 0.05. One-way ANOVA with Dunnett’s test was used for group comparisons, except for Figure 4, which used nested ANOVA with Tukey’s test. Sham/Arrest time points were compared to Naïve and to each other. Analyses were done with GraphPad Prism 8 (San Diego, CA).

## Results

We used a murine model to study physiological, transcriptomic, and immune responses after cardiac arrest. Early changes in EF, mRNA expression, and immune cell populations peaked within the first day, with gradual recovery. We identified specific inflammatory pathways, emphasizing the acute and transient nature of the response.

### Establishment of a murine cardiac arrest model with representative echocardiography and ECG data

Age, sex, and weight-matched C57BL/6J mice were divided into Naïve, Sham, and Arrest groups (Table 1). Naïve mice underwent baseline echocardiography prior to tissue collection. Sham and Arrest mice underwent procedure and were evaluated at either 0.5, 1, 3, or 7 days post-arrest (Figure 1A). Animals in the Arrest group had a mean asystole duration of 8.3 min and total time to extubation of 22.0 min (Table 1). There was no significant drop in body temperature over the period of the experiment. Cardiac EF was significantly lower in Arrest mice at 0.5 days and 1 day post-procedurally when compared to both Naïve and Sham mice at their respective time points (Fig. 1B). No death was recorded in the Naïve or Sham groups. 1 mouse from the 3-day Arrest group and 2 mice from the 7-day Arrest groups died prior to end-point studies (Table 1, Fig. 1C). In the 3-day and 7-day Arrest groups, the differential changes in EF in the CA group was not significant in 3-day or 7-day Arrest mice, signifying gradual recovery of EF as previously reported in this model^18^.

**Table 1.** Physiologic and survivorship data across sample groups. <u>Return of spontaneous circulation</u> <u>(ROSC)</u> indicates the amount of time elapsed between 8-min asystole time and the first observed ventricular complex by ECG and confirmation with Doppler flow analysis. <u>Time to extubation</u> is the total duration of time from procedure start to extubation. Data expressed ± standard deviation. Weights are reported from the day of procedure. Temperatures are reported 1 hour after procedure. *Animal death post-op day 2 (1 female), **Animal death post-op days 1 and 2 (both males).

| Group | <u>Number of Animals Studied (Total)</u> |  |  | <u>Weight (g)</u> |  |  | <u>Temp after procedure (°C)</u> |  | <u>Time to ROSC (min)</u> | <u>Time to extubation (min)</u> |
| --- | --- | --- | --- | --- | --- | --- | --- | --- | --- | --- |
|  | Naïve | Sham | Arrest | Naïve | Sham | Arrest | Sham | Arrest | Arrest | Arrest |
| **0 d** | 5 (5) | n/a | n/a | 23.1±3.1 | n/a | n/a | n/a | n/a | n/a | n/a |
| **0.5 d** | n/a | 4 (4) | 4 (4) | n/a | 24.2±3.4 | 21.7±1.9 | 36.7±0.4 | 36.7±0.5 | 0.22±0.05 | 21.4±1.9 |
| **1 d** | n/a | 4 (4) | 4 (4) | n/a | 23.0±3.2 | 22.8±3.3 | 36.5±0.2 | 36.8±0.5 | 0.50±0.16 | 23.8±1.6 |
| **3 d** | n/a | 4 (4) | 4 (5)* | n/a | 22.1±2.8 | 21.7±3.1 | 36.8±0.1 | 36.6±0.2 | 0.26±0.05 | 22.3±1.3 |
| **7 d** | n/a | 4 (4) | 4 (6)** | n/a | 22.4±4.3 | 21.8±2.8 | 36.7±0.1 | 36.1±0.2 | 0.23±0.03 | 21.0±1.6 |

### Transcriptomic Profiling Revealing Early and Transient Pathway Alterations Following Cardiac Arrest in Murine Models

Next, we conducted a bulk RNA sequencing study to identify and characterize the molecular changes occurring in the heart following CA, providing insight into the gene expression pathways involved in the early post-arrest response. Understanding these transcriptomic alterations is crucial for uncovering potential therapeutic targets and improving outcomes in cardiac arrest recovery. Bulk RNA sequencing was performed on homogenized left ventricles from each treatment group at multiple time points. Differentially expressed genes were identified by comparing Naïve mice to both Sham and Arrest groups (Supplemental Figure 1). Principal component analysis (PCA) of the data was conducted to compare transcriptomic profiles across Naïve, Sham, and Arrest groups at various time points (0.5, 1-, 3-, and 7-days post-procedure; Figure 2A). The most pronounced transcriptomic changes occurred at 0.5 and 1 day in both Sham and Arrest mice, with these alterations normalizing by 3 and 7 days.

**Figure 2.**
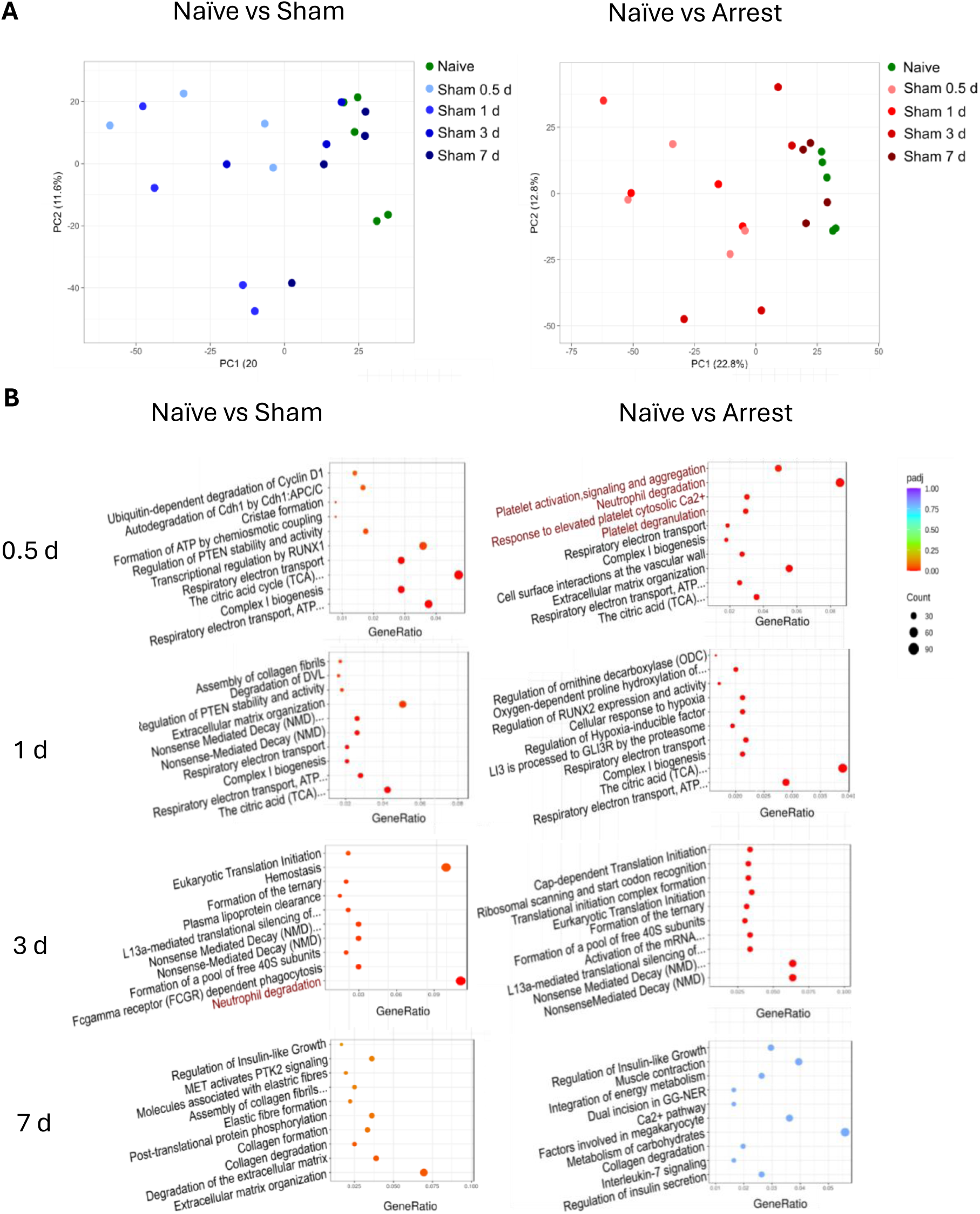
Transcriptomic comparison and pathway Analysis of Sham and Arrest myocardial changes to Naïve controls. A. Principal component analysis (PCA) plots of Naïve transcriptomes compared to Sham (left) or Arrest (right) at 0.5, 1, 3, and 7 d, demonstrating resolution of changes over time. B. Top 10 Reactome pathway analyses comparing Naïve controls to Sham (left) and Arrest (right) hearts at study end-points. Inflammation associated pathways are highlighted in red.

Reactome pathway analysis was utilized to identify broad categories of transcriptome changes (Figure 2B). The top 10 Reactome pathways are presented for each time point by adjusted p-value, allowing comparison of Naïve controls to Sham and Arrest animals. The most prominent pathways in both the Sham (who receive anesthesia & epinephrine) and Arrest (anesthesia & epinephrine & asystole) groups predominantly involve mitochondrial and energetic pathways at 0.5- and 1-day (e.g., citric acid cycle, respiratory electron transport, complex I biogenesis). The Arrest mice, however, feature a number of inflammatory pathways (e.g. platelet activation, neutrophil degranulation) at 0.5 days, which are not present at later time points. Of note, the Sham groups do contain some inflammation associated pathways (e.g. neutrophil degranulation) at the 1-d end-point, though these pathways do not fall into the top ten change (see Supplemental Table 3 for full list of significant changes). By 7 days, there were minimal significant pathway changes in either group. No pathways were significantly different between Naïve and Arrest mice at 7 days, while only three pathways (related to extracellular matrix and collagen degradation) remained significant between Sham and Naïve mice at this time point, suggesting a near-complete resolution of transcriptomic changes. Similar pathway comparisons between Sham and Arrest groups revealed only a few significant alterations, most notably involving extracellular matrix organization at 0.5 and 1 day (Supplemental Figure 2).

### Immune Profiling Reveals Dynamic Inflammatory Responses Following Cardiac Arrest

In order to more deeply characterize the inflammatory changes implicated by RNA sequencing, immunoprofiling was completed on transcriptome data to assess specific immune cell groups that may be contributing to bulk RNA changes. 16 immune cell populations were assessed by comparing gene profiles linked to specific populations and generating the z-scores of mRNA expression changes between all treatment groups (Figure 3A). Heat maps of these z-scores indicate a marked upregulation of immune cell-related genes, most prominently at 1 day in Sham mice compared to Naïve, and at 0.5 days in Arrest mice compared to Naïve. Notably, significant upregulation of neutrophils and monocyte-associated genes was observed at 0.5 days in Arrest mice and at 1 day in Sham mice (Figure 3B). Additionally, dendritic cells (DCs) and T Helper 2 (Th2) cells were elevated in Arrest mice at 0.5 days compared to Naïve animals, with no other immune cell groups showing significant elevations in Sham or Arrest mice at other time points.

**Figure 3.**
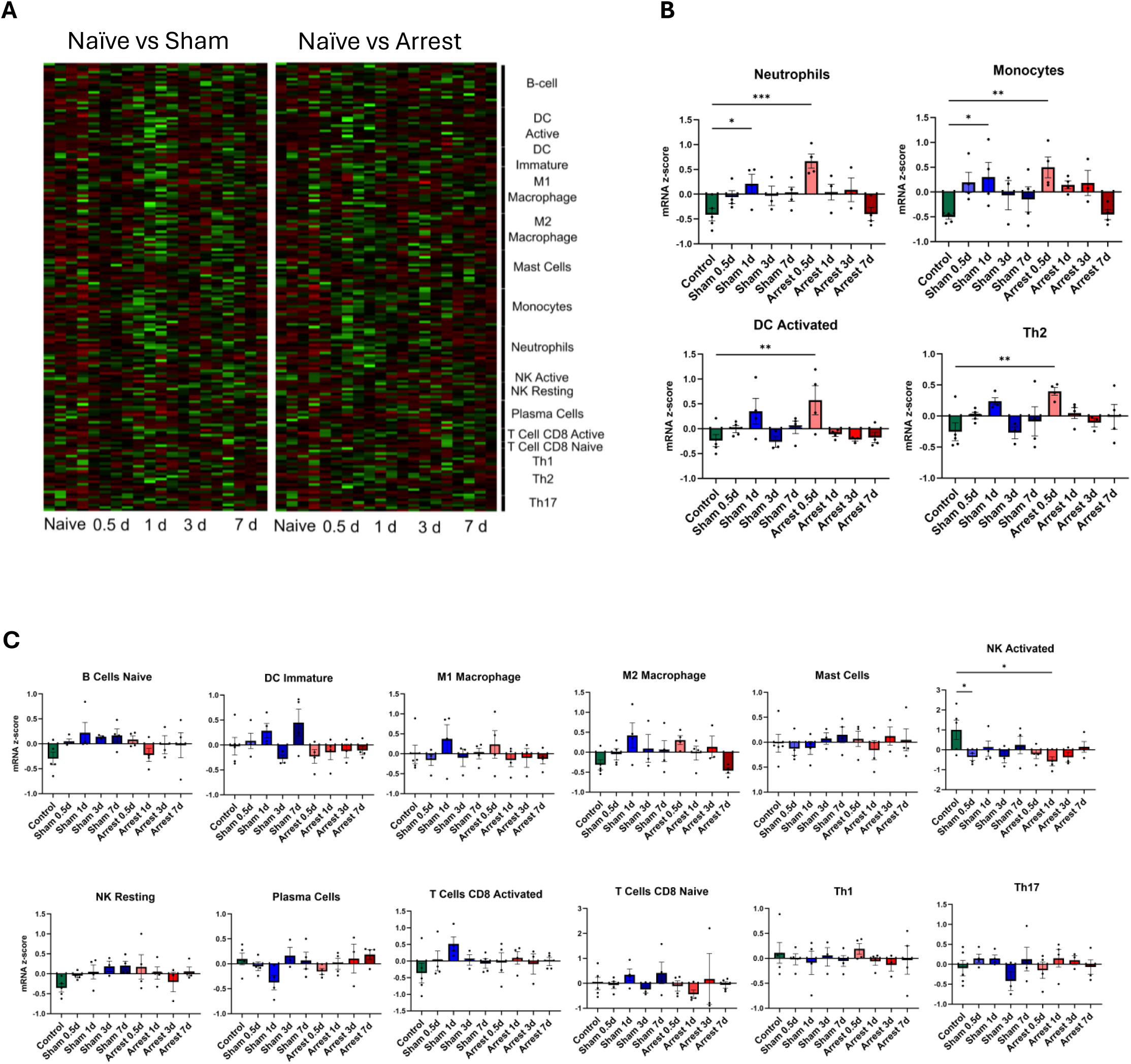
Immunoprofiling of Sham and Arrest hearts based on transcriptome data. A. Heat maps demonstrating up- (green) or downregulation (red) of genes associated with 16 immune cell populations types based on z-scoring of gene expression for Sham (left) and Arrest (right) mice compared to Naïve controls. B. Comparison of average z-score change for neutrophils, monocytes, activated dendritic cells (DC), and T-helper 2 cells (Th2) comparing Sham and Arrest end-points to Naïve controls. C. Comparison of remaining immune cell populations between end-points and Naïve controls. n=4-5/group. *=p <0.05, **=p<0.01, ***=p<0.001 by ANOVA with Dunnett’s multiple comparison test.

To validate and further characterize immune cell infiltration into the myocardium, we conducted flow cytometry on washed and digested ventricular walls from Naïve, Sham, and Arrest mice at each time point. Immune cell identification via flow cytometry was confirmed using spleen tissue with single-label controls (Supplemental Figure 3). CD45+ and CD3+ cells were significantly elevated in Arrest mice at 0.5 days compared to Naïve (Figure 4A). Ly6G+ CD11b+ neutrophils were also elevated in Arrest mice at 0.5 days compared to both Naïve and Sham mice. Sham mice exhibited an increased concentration of macrophages at 1 day compared to Arrest mice at the same time point with elevated M1 macrophages, specifically. 1-day Sham mice also showed elevated DCs compared to Arrest mice at the same time point, and finally, natural killer (NK) cells were significantly elevated in 3-day Sham mice when compared to both Naïve and 3-day Arrest mice.

**Figure 4.**
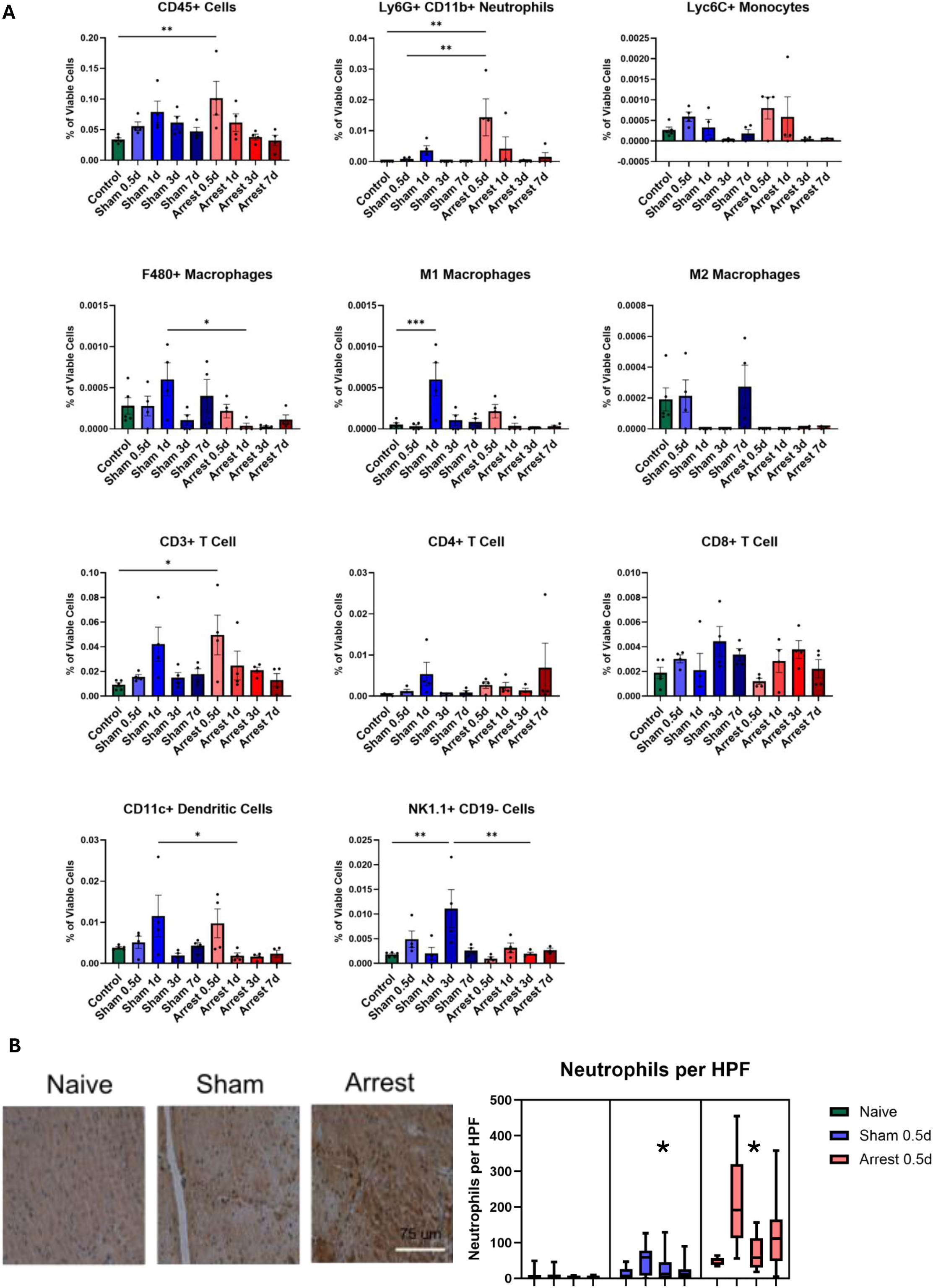
Quantification of immune cell populations in Sham and Arrest hearts by flow cytometry. A. Quantities represent the percent of viable cells that stained positive for markers of specific immune population. Comparisons are made between all end-points to Naïve controls as well as between same day Sham and Arrest end-points n=4-5/group. *=p <0.05, **=p<0.01, ***=p<0.001 by ANOVA with Dunnett’s multiple comparison test. B. Representative histologic sections with DAB staining of myeloperoxidase as a marker of neutrophil infiltration in Naïve controls as well as Sham and Arrest hearts from 0.5 d. n=4/group. *=p<0.05 compared to Naïve controls by nested 1 way ANOVA with Tukey’s multiple comparison test.

### Histological Confirmation of Increased Neutrophil Infiltration in the Cardiac Tissues of Arrest Mice at 0.5 Days Post-Procedure

Next, we wanted to confirm the presence of infiltrated neutrophils in the affected cardiac tissues in our CA model. We performed histologic staining to confirm the changes in infiltrated neutrophil numbers observed in the Arrest group at 0.5 days. Analysis revealed a significantly higher neutrophil count per high-powered field in the cardiac tissue of Arrest mice compared to both Naïve and Sham mice (Fig. 4B). This increased neutrophil presence aligns with the Immunoprofiling data, further validating the acute inflammatory response observed shortly after cardiac arrest.

### Flow Cytometry Analysis revealed Modulation of Natural Killer Cells but not of other immune cells in the Peripheral Blood among the Experimental Groups

Flow cytometry analysis was performed on peripheral blood mononuclear cells (PBMCs) across all treatment groups to assess systemic immune cell alterations following cardiac arrest. Notably, there was a significant decrease in natural killer (NK) cells in the Arrest mice at 1 day post-procedure compared to Naïve mice, indicating a potential suppression or redistribution of these cells in response to CA (Figure 5). Despite this change, the overall infiltration of other immune cell populations remained consistent across the experimental groups, suggesting that the observed inflammatory response is primarily localized to specific immune cell types, such as NK cells, without widespread alterations in other circulating immune cells.

**Figure 5.**
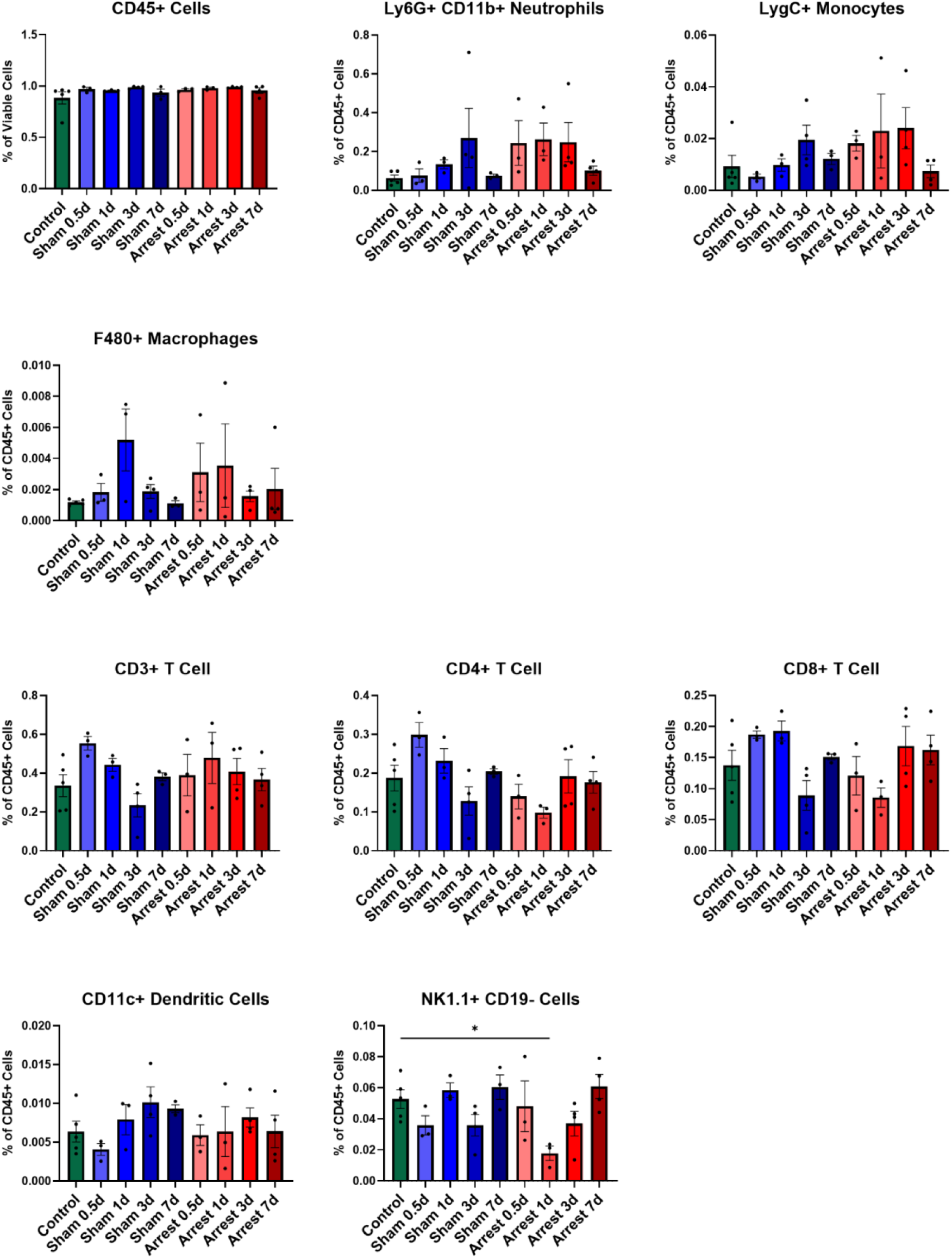
Quantification of immune cell populations in Sham and Arrest peripheral blood by flow cytometry. Quantities represent the percent of viable cells that stained positive for markers of specific immune population. Comparisons are made between all end-points to Naïve controls as well as between same day Sham and Arrest end-points. A full gating strategy is available in Supplementary Figures 3-4. n=3-5/group. *=p <0.05 by ANOVA with Dunnett’s multiple comparison test.

### Elevation of Pro-Inflammatory Cytokine Signaling in the Acute Phase in Response to Cardiac Arrest

To investigate the systemic inflammatory response following cardiac arrest, we conducted a comprehensive cytokine profiling analysis on plasma samples collected at various time points from all treatment groups. The analysis revealed a significant increase in the concentrations of pro-inflammatory cytokines IL-6 and TNF-α in the Arrest mice at 0.5 days post-procedure compared to Naïve controls, indicating an acute inflammatory response shortly after the cardiac event. Additionally, the Arrest group at 3 days post-procedure exhibited elevated levels of IL-1β, suggesting a sustained but evolving inflammatory response. However, no other cytokines showed significant changes between the groups at any of the time points examined, highlighting the specificity of these pro-inflammatory cytokines in the early stages of post-arrest inflammation. This cytokine profile underscores the role of IL-6, TNF-α, and IL-1β as key mediators of the peripheral inflammatory response following cardiac arrest.

## Discussion

This study offers a detailed analysis of physiological, transcriptomic, and immunometabolic responses to cardiac arrest (CA) in a mouse model, revealing key pathophysiological insights. Our findings highlight the rapid onset of inflammation, early gene expression changes, and dynamic immune cell activity in the heart post-arrest. These observations suggest potential therapeutic targets to improve clinical outcomes. Recent trials targeting inflammation post-CA have shown promise with steroids and anti-cytokine therapies^24–26^. We traced transcriptomic and inflammatory changes over time using a controlled mouse model, which minimizes local injury for clearer insights. Transcriptomics revealed significant activation of mitochondrial and inflammatory pathways shortly after CA, normalizing by day 7, suggesting a brief but intense response.

In our study, the significant reduction in cardiac EF observed within the first day post-arrest is indicative of acute myocardial dysfunction, commonly known as myocardial stunning, which typically follows ischemic events (Fig. 1B). The rapid decline in EF, followed by a gradual recovery, aligns with previous studies, highlighting the transient nature of post-CA myocardial dysfunction^13^. Here, we report an absence of significant EF differences at 3- and 7-days post-arrest, which suggests a recovery process, although the molecular mechanisms driving this recovery require further investigation. These observations underscore the importance of early intervention and the potential for therapies that can mitigate the initial decline in cardiac function. Our bulk RNA sequencing revealed significant transcriptomic changes in the myocardium, particularly at 0.5- and 1-day post-arrest. The pronounced activation of mitochondrial and inflammatory pathways during this period suggests a rapid cellular response to ischemic injury and subsequent reperfusion^27^. The early activation of pathways related to energy metabolism and immune responses likely reflects the myocardium’s attempt to restore homeostasis and repair cellular damage. By 3- and 7-days post-arrest, the normalization of these transcriptomic changes indicates a resolution phase (Figure 2), suggesting either a return to a pre-injury state or the establishment of a new equilibrium following the acute phase of injury. Interestingly, although both the Sham and Arrest groups initially exhibited similar transcriptomic changes, the inflammatory response was more pronounced in the Arrest group, particularly at 0.5 days post-arrest. This suggests that while either the surgical procedures or epinephrine treatment (as represented by the Sham group) can induce a mild inflammatory response, the ischemia/reperfusion insult associated with CA exacerbates this response, particularly in pathways involving neutrophil activity and antigen processing. The resolution of these changes by day 7, in parallel with EF recovery, suggests that the inflammatory response in this model is acute and self-limiting.

Our immunoprofiling and flow cytometry analyses provided crucial insights into immune cell dynamics following CA. Immunoprofiling identified early neutrophil and monocyte infiltration into the heart, supporting the acute inflammatory response. Neutrophil presence peaked at 0.5 days, with dendritic cells and T Helper 2 cells also elevated, indicating a complex immune response. Cytokine analysis showed early spikes in IL-6, TNF-α, and IL-1β, suggesting a targeted pro-inflammatory reaction. The transient nature of the inflammatory response is further supported by the observation that neutrophil infiltration peaks early (at 0.5 days) and then diminishes over time, indicating a rapid mobilization of immune cells to the site of injury, followed by a resolution phase as myocardial tissue begins to recover. The specific increase in dendritic cells (DCs) and T Helper 2 cells in the Arrest group at 0.5 days also points to a complex interplay between immune cell types in orchestrating the inflammatory response (Figure 5). Moreover, systemic immune cell alterations, particularly the decrease in NK cells observed at 1 day post-arrest, may reflect a redistribution of immune cells or a transient suppression of certain immune responses in the context of CA. This finding raises interesting questions about the broader systemic effects of CA on immune function and suggests potential areas for further research.

Our cytokine profiling in the CA models revealed elevated levels of IL-6, TNF-α, and IL-1β in the early stages post-arrest, indicating a robust pro-inflammatory response. These cytokines, known mediators of inflammation, have been implicated in the pathogenesis of various cardiovascular conditions, including myocardial infarction and heart failure (Figure 6). The early spike in IL-6 and TNF-α, followed by sustained IL-1β elevation at 3 days, suggests a dynamic inflammatory response that evolves over time. The specificity of these cytokine changes, with no significant alterations in other cytokines, underscores the targeted nature of the immune response following CA. This specificity may have important implications for developing therapeutic strategies that modulate the inflammatory response without broadly suppressing immune function, which could be detrimental.

**Figure 6.**
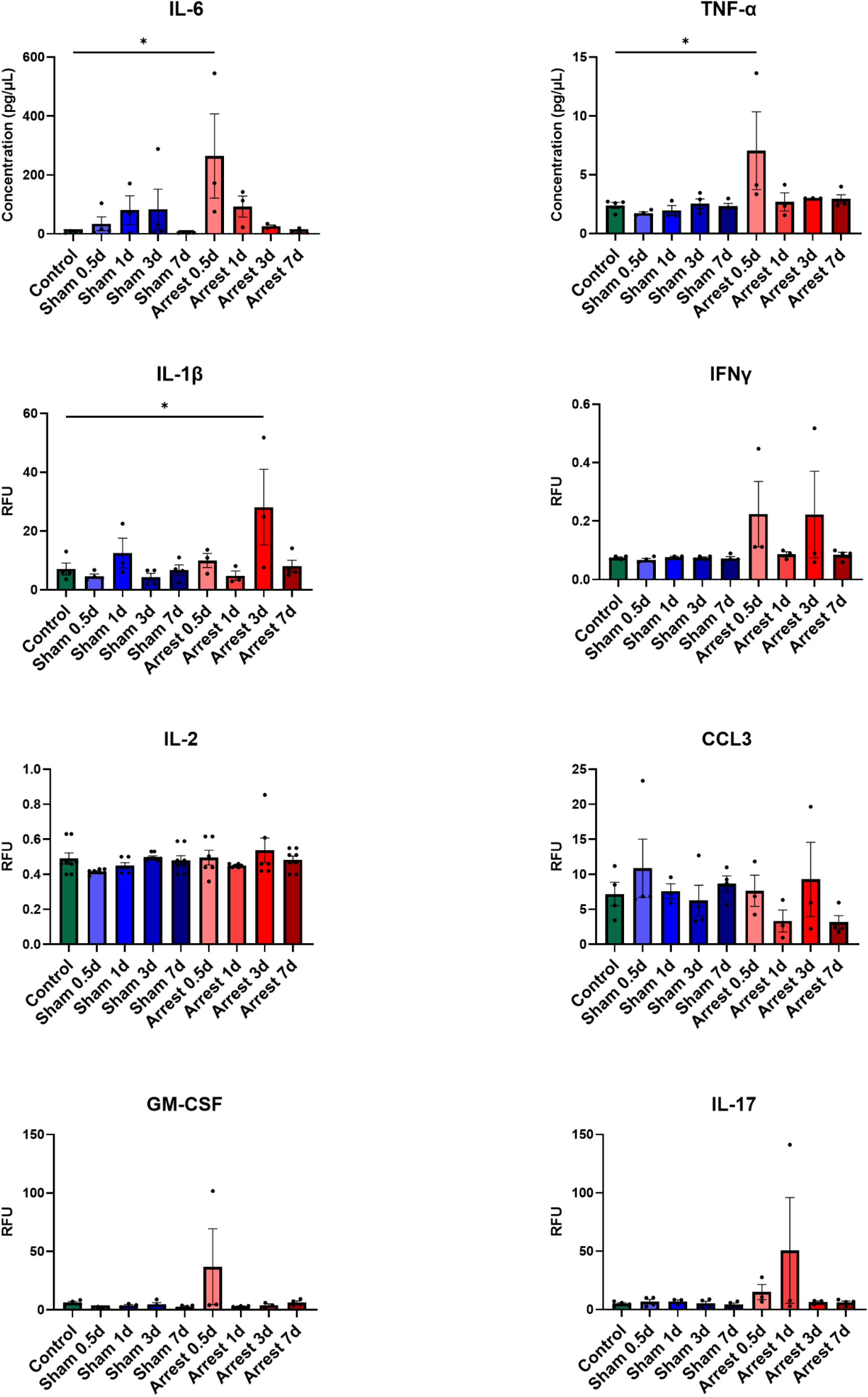
Peripheral blood cytokine levels. Blood from each end-point was evaluated for cytokine levels by multiplex assay kit for levels of interleukin 6 (IL-6), tumor necrosis factor α (TNF- α), interleukin 1β (IL1 β), interferon γ (IFN γ), interleukin 2 (IL2), chemokine ligand 3 (CCL3), granulocyte-macrophage colony-stimulating factor (GM-CSF), and interleukin 17 (IL-17). n=3-5/group. *=p <0.05 by ANOVA with Dunnett’s multiple comparison test.

These results underscore the potential for therapies that target specific pathways to mitigate damage and promote recovery. However, translation to clinical practice will require further validation in humans and safe therapeutic strategies.

## Limitations of this Study

This study has several limitations. The use of young, healthy mice without underlying cardiac disease may not fully represent the clinical population of CA patients. Furthermore, we did not examine hypoxia-induced brain damage nor did we assess the extent of myocardial damage and necrosis following CA, though previous work in this model has found no cardiomyocyte death^28^. In reported models of myocardial stunning, early studies identified neutrophils as significant contributors to ventricular wall dysfunction^29^, however, subsequent studies involving neutrophil depletion in a pig model of myocardial stunning did not show improvement in cardiac dysfunction^30^. The efficacy of systemic anti-neutrophil therapies in human cardiac arrest patients remains unexplored. Notably, our study did not find a strong signal for macrophage activation. Although circulating cytokine levels, including TNF-α and IL-6, were acutely elevated in the CA model, we did not observe an increase in circulating immune cells in plasma samples. The Sham model, which involved epinephrine treatment and surgical intervention, also triggered an inflammatory response, peaking around one day after the insult. Previous studies have shown that epinephrine alone can cause immune activation and neuroimmune communication^31^, and these pro-inflammatory effects of epinephrine therapy suggest a limited role for epinephrine in resuscitation efforts^32^.

In conclusion, understanding the acute inflammatory response in CA can inform targeted interventions, potentially reducing high morbidity and mortality rates. Future research should explore these pathways and test new treatments to enhance recovery and survival.

## Acknowledgments

CR was responsible for the conceptualization of these studies. CR designed the methodology, and with SP, AJ, AE, ST, and DR performed the investigation. Formal analyses were performed by SP, ST, and CR. SP and CR completed visualization. SP, KR, and CR wrote the manuscript. CR supervised the project and provided resources for its completion. All authors reviewed the final manuscript and are responsible for its integrity.

## Abbreviations

ACK: Ammonium-Chloride Potassium lysis buffer
CA: Cardiac Arrest
CCL3: Chemokine ligand 3
DAB: 3,3’-diaminobenzidine
DC: Dendritic Cell
EF: Ejection Fraction
ELISA: Enzyme linked immunosorbent assay
FACS: Fluorescence-activated Cell Sorting
GM-CSF: Granulocyte-macrophage colony-stimulating factor
HBSS: Hanks Balanced Salt Solution
HPF: High Powered Field
IL: Interleukin
LV: Left Ventricle
MI: Myocardial Infarction
MPO: Myeloperoxidase
NK: Natural Killer cell
PBMC: Peripheral blood mononuclear cells
PCAS: Post-Cardiac Arrest Syndrome
ROSC: Return of Spontaneous Circulation

## Figure Legends

**Supplemental Figure 1. Volcano plots of differentially expressed genes between groups.** All Sham and Arrest end-points are compared to a single group of Naïve controls. Sham vs Arrest groups are compared at the same endpoint (i.e. 0.5 d sham vs 0.5 d arrest).

**Supplemental Figure 2. Pathway Analysis of Sham and Arrest myocardial at each end-point.** The top 10 Reactome pathway analyses comparing Sham and Arrest hearts at study end-points.

**Supplemental Figure 3. Gating strategy for infiltrating immune cells taken from a representative spleen.**

**Supplemental Figure 4. Gating strategy for infiltrating immune cells taken from a representative left ventricle.**

**Supplemental Figure 5. Gating strategy for immune cells taken from peripheral mononuclear blood cells.**

