## Supplemental Figures for "Temporal dynamics of inflammatory and transcriptome changes in the myocardium in a murine model of cardiac arrest"

Supplementary Figure 1

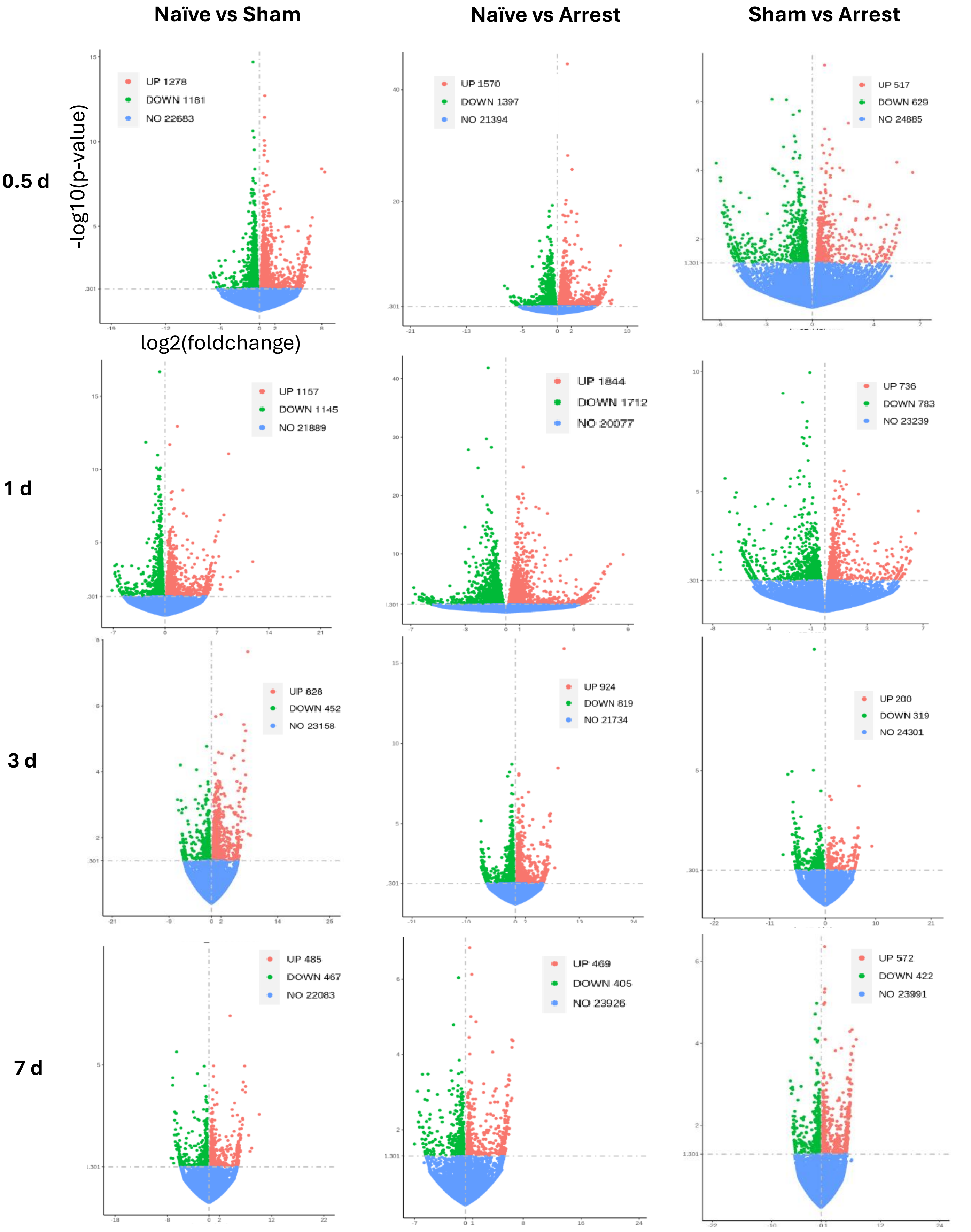

Supplementary Figure 2

### Sham vs Arrest Reactome Pathways

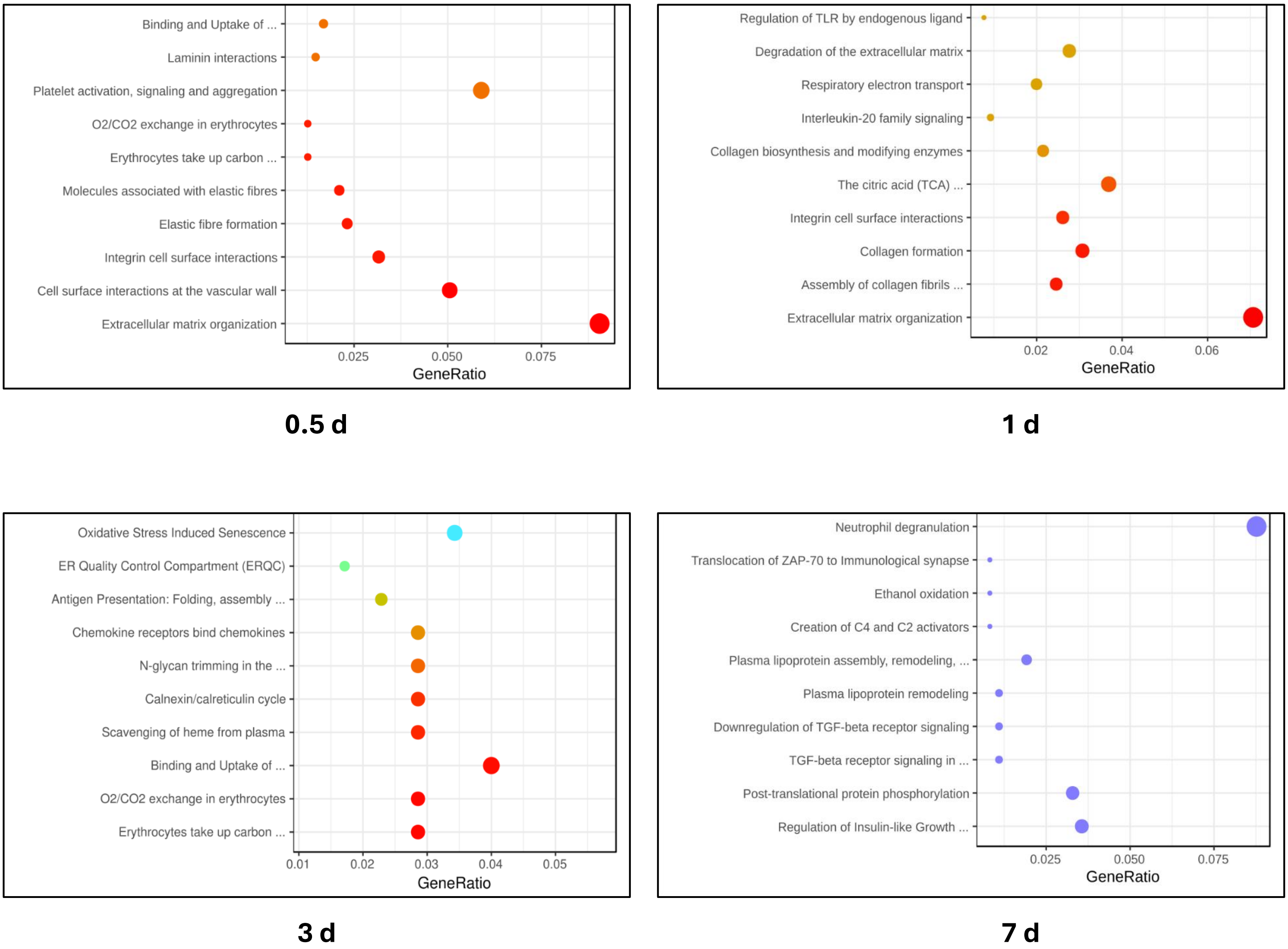

Supplementary Figure 3

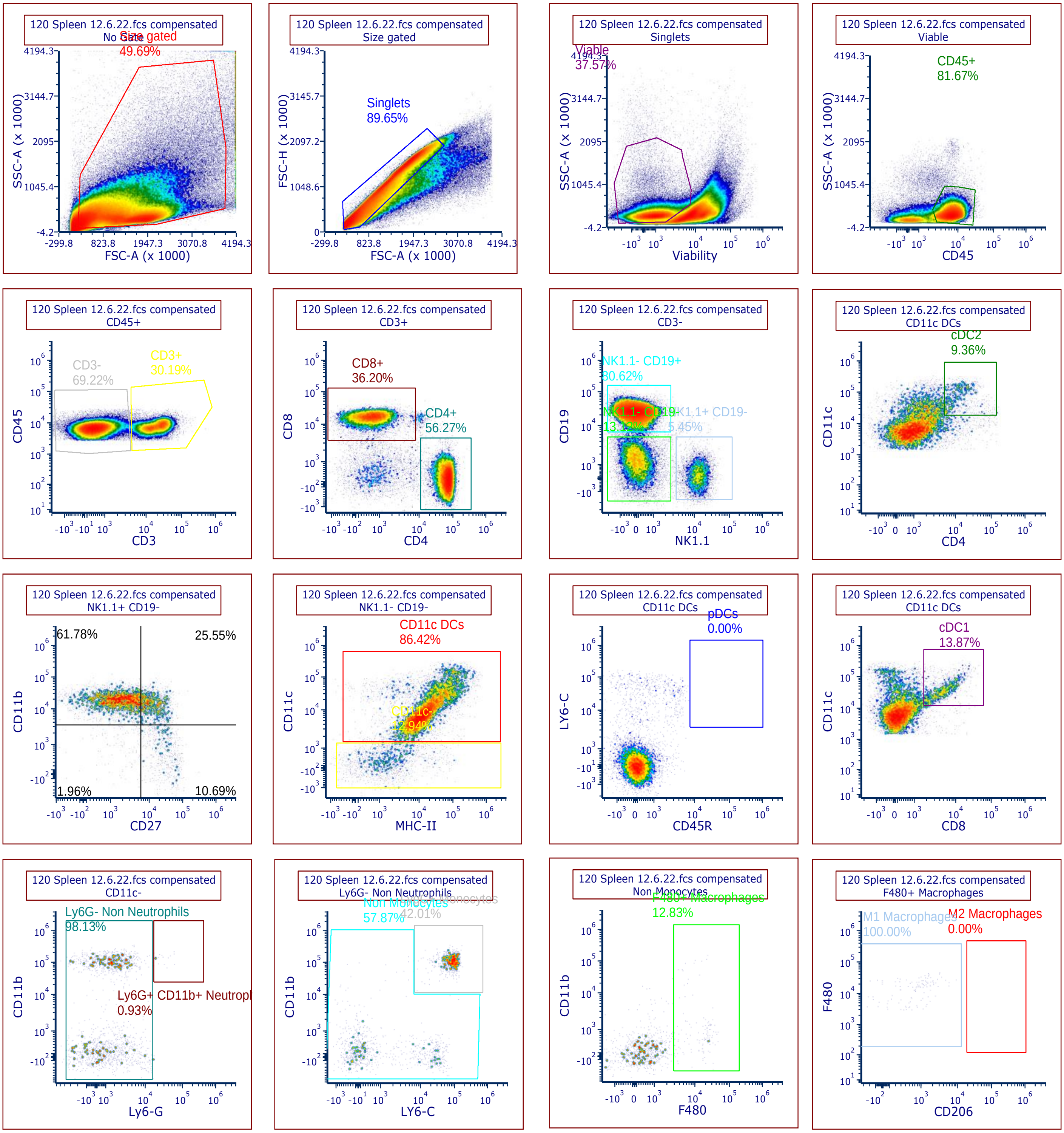

Supplementary Figure 4

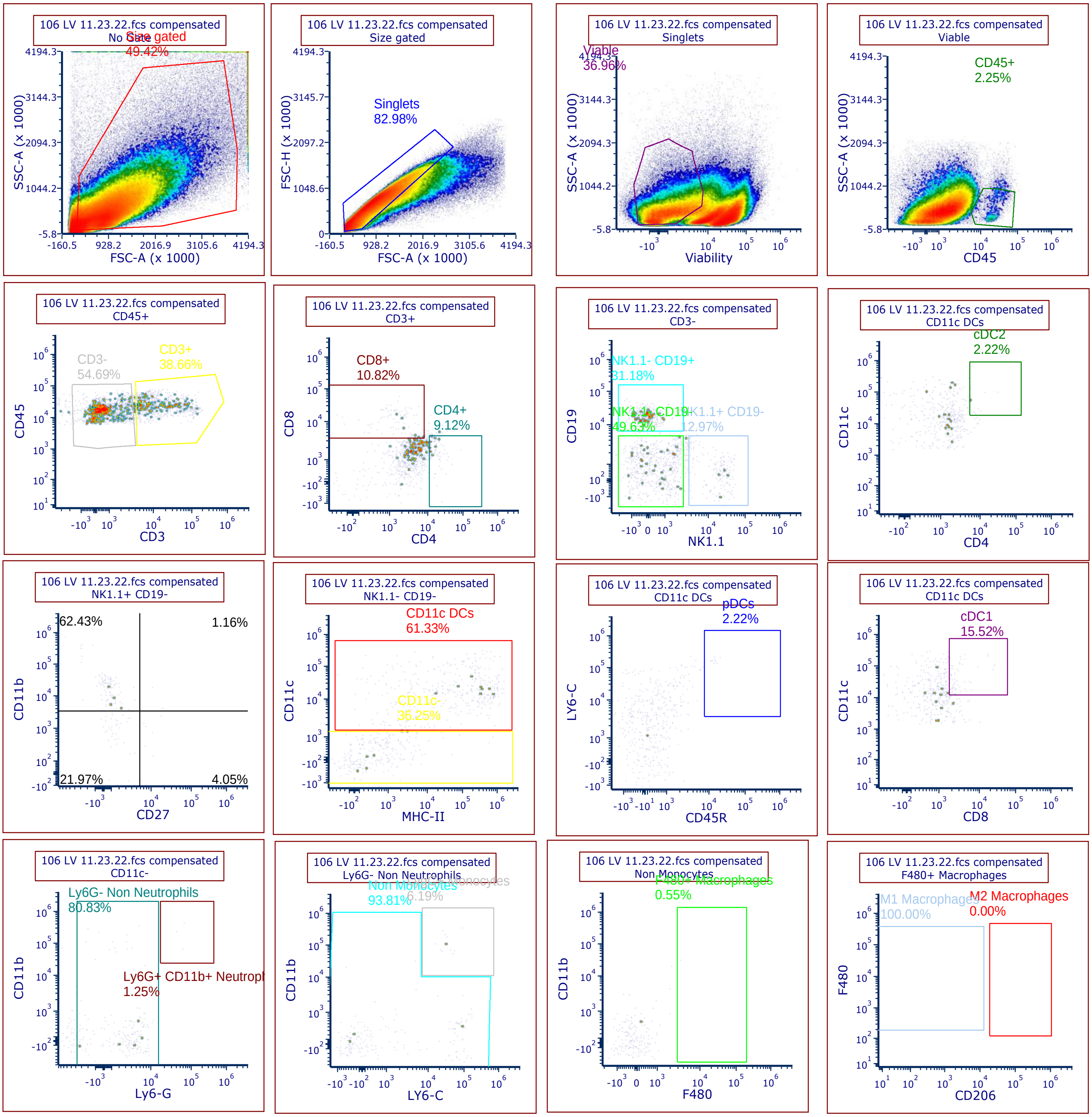

Supplementary Figure 5

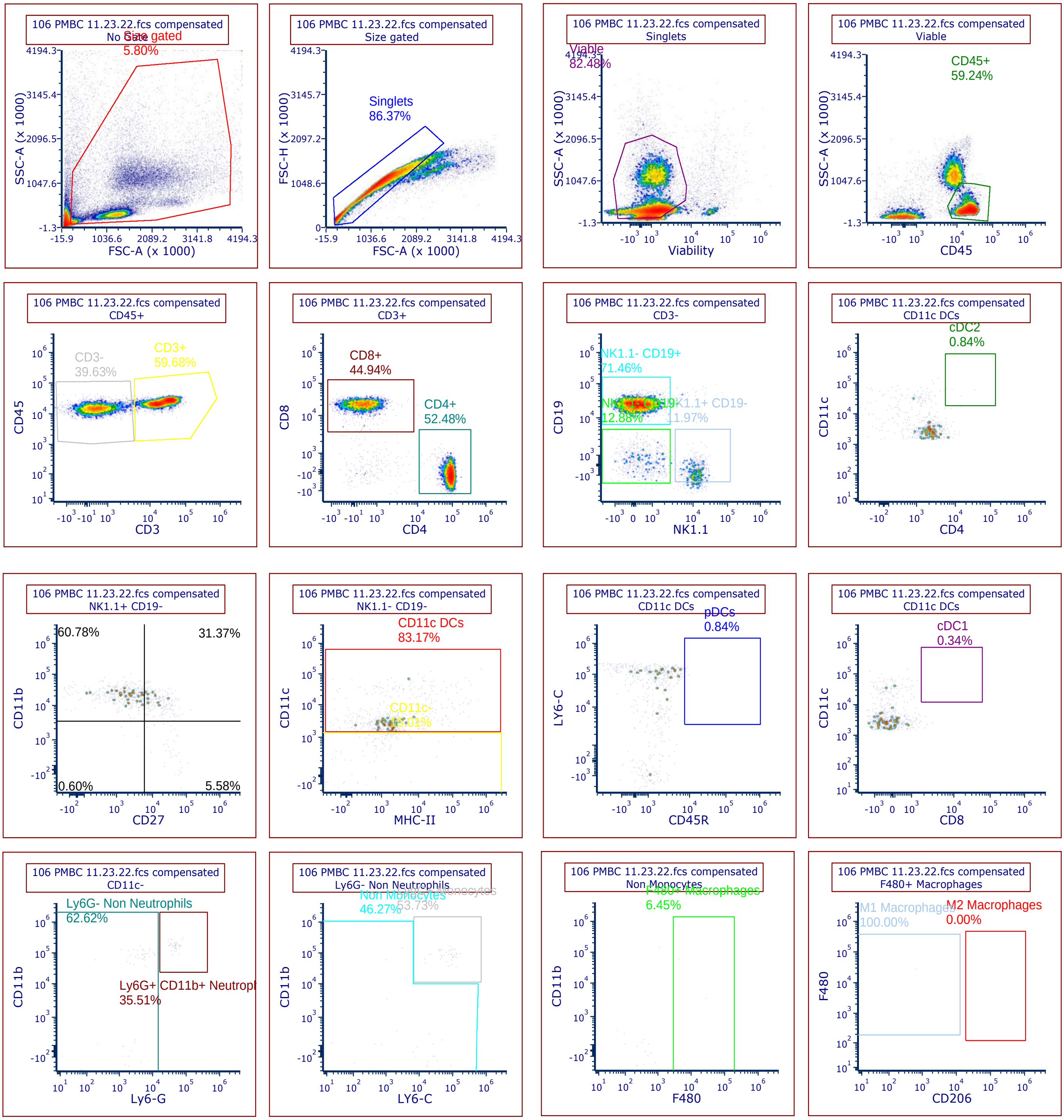
